# Gut microbiota differently contributes to intestinal immune phenotype and systemic autoimmune progression in female and male lupus-prone mice

**DOI:** 10.1101/787440

**Authors:** Benjamin M. Johnson, Marie-Claude Gaudreau, Radhika Gudi, Robert Brown, Gary Gilkeson, Chenthamarakshan Vasu

**Affiliations:** Microbiology and Immunology, College of Medicine, Medical University of South Carolina, Charleston, SC-29425; Division of Rheumatology, Department of Medicine, College of Medicine, Medical University of South Carolina, Charleston, SC-29425

**Keywords:** systemic lupus erythematosus, gut mucosa, gut microbiota, testosterone, autoimmunity, gender bias, inflammation

## Abstract

The risk of developing systemic lupus erythematosus (SLE) is about 9 times higher among women compared to men. Our recent report, which used (SWRxNZB) F1 (SNF1) mouse model of spontaneous lupus, showed a potential link between immune response initiated in the gut mucosa at juvenile age (sex hormone independent) and SLE susceptibility. Here, this mouse model, we show that gut microbiota contributes to a pro-inflammatory immune response in the intestine and autoimmune progression, primarily in females, leading to an associated gender bias. We found that gut microbiota composition in male and female littermates are significantly different only at adult age and depletion of gut microbes causes suppression of autoimmune progression only in females. In agreement, microbiota depletion suppressed the pro-inflammatory cytokine response of gut mucosa in both juvenile and adult females. In male SNF1 mice, on the other hand, orchidectomy (castration) caused changes in the composition of gut microbiota and a modest acceleration of autoimmune progression. However, cecum microbiota transplantation studies failed to show superior protection of females from autoimmunity by androgen-influenced gut microbiota. Overall, our work shows that microbiota-dependent pro-inflammatory immune response in the gut mucosa of females initiated at juvenile ages and androgen-dependent protection of males contributes to gender differences in the intestinal immune phenotype and systemic autoimmune progression.

## 2. Introduction

Systemic lupus erythematosus (SLE), an autoimmune disease, arises when abnormally functioning B lymphocytes produce auto-antibodies to DNA and nuclear proteins, resulting in immune complexes that cause damage to the tissue [1]. While the triggers are not known, it is generally believed that SLE is caused by a combination of genetic and environmental factors [1–3]. Dietary factors can also have a profound impact on the gut and systemic immune responses [4–8]. Recent studies that used human samples and rodent models have shown that gut microbiota composition influences the rate of disease progression and the overall disease outcome [9–13]. In addition to direct effects on the systemic and gut immune cell functions, dietary factors can change the composition of gut microbiota and affect autoimmune outcomes [10, 14, 15]. Recent reports from our laboratory demonstrated that minor dietary deviations such as changes in the pH of drinking water alter the composition of gut microbiota and incidence of autoimmune diseases including SLE [13, 16].

Importantly, there is a predisposition for SLE in women with a prevalence ratio of about 9:1 over men [17–19]. Numerous studies have suggested that sex hormones, estrogen in particular, can contribute to the onset and development of disease activities associated with SLE. Estrogen is reported to have inductive effects on autoimmune-related immune responses such as production of antibody and pro-inflammatory cytokines [17, 19–21]. Studies using the non-obese diabetic (NOD) mouse model of T1D have shown a role for testosterone-influenced microbiota, independent of estrogen-mediated effects, in determining the gender bias of autoimmune diseases [22, 23]. These reports suggested a strong involvement of gut microbial factors in determining gender disparity in systemic autoimmunity. Although it has been shown that lupus-susceptible GF mice develop disease [24, 25], it is not known if microbiota affects gut immune phenotype and modulates disease outcomes differently in males and females under lupus susceptibility. Further the contribution of testosterone influenced microbiota, as reported in NOD mice [22, 23], to gender bias in lupus autoimmunity is largely unknown. Although MRL/lpr mice showed gender specific differences in gut microbiota close to disease onset [26], it is not known if these differences are hormone driven, or can impact intestinal immune phenotype and autoimmune progression.

Our recent study that used lupus-prone (SWRxNZB)F1 (SNF1) mice, for the first time, showed the potential contribution of immune response initiated in the gut mucosa in lupus associated gender bias [27]. We also showed that pro-inflammatory immune response in the gut mucosa of lupus-prone females can be detected as early as at juvenile age, an age at which there is no sex hormone produced and much before the appearance of systemic autoimmunity in these mice. In another report, we showed that change in the pH of drinking water provided to this mouse model has a profound effect not only on the composition of gut microbiota and the type of intestinal immune response, but also on the disease progression [13]. These observations suggest a potential role for gut microbiota composition and mucosal immunity in shaping the gender bias of systemic autoimmunity in lupus-prone background.

Here we show that gut microbiota composition in lupus-prone SNF1 mouse littermates, are different at adult ages, but not at juvenile age. Interestingly, depletion of gut microbiota starting at juvenile age suppressed the inflammatory phenotype, ameliorated systemic autoantibody production and disease incidence only in females. On the other hand, castration of male SNF1 mice resulted in altered gut microbiota and modest increases in autoantibody production and lupus incidence, as compared to microbiota from females and castrated males. However, androgen influenced gut microbiota did not show better ability to alter the disease outcomes in female mice. Overall, these observations show that juvenile age onset and progressive increase in intestinal pro-inflammatory immune response, and adult age systemic autoantibody production in lupus-prone females are gut mucosa-microbiota interaction dependent. Further, androgen, microbiota independently, promotes protection of lupus-prone males from autoimmunity at adult ages and contributes to gender bias in lupus.

## 3. Materials and Methods

### 3.1. Mice

SWR, NZB mice, and C57BL/6 (B6) mice were purchased from the Jackson Laboratory (Bar Harbor, Maine) and housed under SPF conditions at the animal facilities of Medical University of South Carolina (MUSC). (SWRxNZB)F1 (SNF1) hybrids were generated at the SPF facility of MUSC by crossing SWR females with NZB males. In some experiments, male SNF1 mice were subjected to orchidectomy (castration) or mock surgery after anesthesia with ketamine and xylozine combination. In some experiments, mice were given a cocktail of broad spectrum antibiotics as described in our recent report [28] to deplete gut microbiota. Depletion of gut microbiota was confirmed by culture of fecal pellet suspension on brain heart infusion agar plates under aerobic and anaerobic conditions as described before [28]. Urine and tail vein blood samples were collected at different time-points to detect proteinuria and autoantibodies. All animal experiments were performed according to ethical principles and guidelines were approved by the institutional animal care committee of MUSC.

### 3.2. Proteinuria

Urine samples collected weekly were tested for protein levels by Bradford assay (BioRad) against bovine serum albumin standards. Proteinuria was scored as follows; 0: 0-1mg/ml, 1: 1-2mg/ml, 2: 2-5 mg/ml, 3: 5-10mg/ml and 4: ≥ 10mg/ml. Mice that showed proteinuria >5 mg/ml were considered to have severe nephritis.

### 3.3. ELISA

Levels of antibodies against nucleohistone and dsDNA in mouse sera were determined by ELISA as described in our recent reports [13, 27]. Briefly, 1.0μg/well of nucleohistone (Sigma-Aldrich) or dsDNA from calf thymus (Sigma-Aldrich) was coated as antigen, overnight, onto ELISA plate wells. Serial dilutions of the sera were made and total IgG and IgG isotypes against these antigens were detected using HRP-conjugated respective anti-mouse immunoglobulin isotype antibodies (Sigma-Aldrich, eBioscience and Invitrogen).

### 3.4. Quantitative PCR

RNA was extracted from 2-cm pieces of the distal ileum or 1cm pieces of distal colon using Isol-RNA Lysis Reagent (5prime) according to manufacturer’s instructions. cDNA was prepared from RNA using first strand synthesis kit (Invitrogen) and PCR was performed using SYBR-green master-mix (BioRad) and target-specific custom-made primer sets. A CFX96 Touch real time PCR machine (BioRad) was used and relative expression of each factor was calculated by the 2Δ-CT cycle threshold method against β-actin control.

### 3.5. 16S rRNA gene targeted sequencing and bacterial community profiling

Total DNA was prepared from fresh fecal pellets of individual mice for bacterial community profiling as described in our recent reports [13, 16, 28]. Briefly, DNA in samples was amplified by PCR using 16S rRNA gene-targeted primer sets to assess the bacterial levels. V3-V4 region of 16S rRNA gene sequencing was performed using Illumina MiSeq platform at MUSC genomic center. The sequencing reads were fed in to the Metagenomics application of BaseSpace (Illumina) for performing taxonomic classification using an Illumina-curated version of the GreenGenes taxonomic database which provided raw classification output at multiple taxonomic levels. The sequences were also fed into QIIME open reference operational taxonomic units (OTU) picking pipeline [29] using QIIME pre-processing application of BaseSpace. The OTUs were compiled to different taxonomical levels based upon the percentage identity to GreenGenes reference sequences (i.e. >97% identity) and the percentage values of sequences within each sample that map to respective taxonomical levels were calculated. The OTUs were also normalized and used for metagenomes prediction of Kyoto Encyclopedia of Genes and Genomes (KEGG) orthologs employing PICRUSt as described before [26, 30–32]. The predictions were summarized to multiple levels and functional categories as well as microbial communities at different taxonomical levels were compared among different groups. Statistical Analysis of Metagenomic Profile Package (STAMP) [31] or web-based MicrobomeAnalyst application [33] were employed for analyzing OTU.Biom table and PICRUSt data for visualization and statistical analyses where appropriate.

### 3.6. Microbiota transfer

Transfer of microbiota was performed as described in our recent reports [13, 16]. Briefly, homogenous suspensions of cecum contents were made in chilled 0.1M bicarbonate buffer and administered to recipients by oral gavage. Suspension from one cecum was given to 5 mice. This process was repeated for three consecutive days using fresh preparations of microbes from additional donors. The recipients were tested for proteinuria and auto-antibody levels as described above.

### 3.8. Multiplex assay / suspension bead array

Immune cells were enriched from collagenase digested intestinal tissues by Percoll gradient centrifugation and culture supernatants were tested for cytokine levels. Luminex multiplex assay based suspension bead array (SBA) was carried out using magnetic bead based cytokine/chemokine 26-plex or IFN-α and IFN-β 2-plex panels from eBiosciences and/or Invitrogen. Assay plates were read and concentrations were calculated using FlexMap3D instrument and xPONENT software (Luminex Corporation) at MUSC’s immune monitoring and discovery core.

### 3.7. Statistical analysis

Proteinuria curves were analyzed using log-rank method (http://bioinf.wehi.edu.au/software/russell/logrank/) and the proteinuria scores were analyzed using Chi-square test. Two-tailed *t*-test or Mann Whitney test was also employed to calculate *p*-values where indicated. A p value of ≤0.05 was considered statistically significant. GraphPad Prism was used for calculating statistical significance. For some 16S rDNA data, Welch’s two-sided t-test was employed and the *p*-values were corrected from multiple tests using the Benjamini and Hochberg approach using STAMP or MicrobiomeAnalyst applications.

## 4. Results

### 4.1. Intestinal immune phenotype in SNF1 mice

SNF1 mice that develop lupus symptoms and proteinuria spontaneously and show gender bias in disease incidence similar to human SLE patients [34, 35]. Recently, we reported that significant amounts of circulating autoantibodies against nucleohistone and dsDNA are detectable in SNF1 mice at 16 weeks of age and severe nephritis indicated by high proteinuria after 22 weeks of age [13, 27]. We also showed that about 80% of the female and 20% of the male SNF1 mice develop severe nephritis within 32 weeks. Here, to further characterize the gut immune phenotype in lupus-prone SNF1 mice, we determined mRNA and secreted protein levels of these key pro-inflammatory cytokines in both large and small intestines of 4or 16 week old (pre-nephritic) male and female littermates. Quantitative PCR analysis shows that most pro-inflammatory factors including *Il6*, *Il4*, *Il9*, *Il17*, *Il21, Ifna and Ifnb* are expressed at higher levels in both colon and ileum of female SNF1 mice as compared to their age-matched male counterparts (Fig. 1A and Supplemental Fig. 1), and lupus-resistant B6 mice (not shown). Significantly higher *Ifng* expression was detected primarily in the large intestine of the SNF females. Further, the expression levels of endosomal TLRs, which are implicated in SLE (*Tlr7*, *Tlr8* and *Tlr9*) [36–40], are also high in both large and small intestines of SNF females. Importantly, significant differences in the expression levels of most pro-inflammatory factors were detectable even in 4 week old (juvenile) mice and a robust increase in the expression levels of these factors in the small and large intestines were detected at 16 weeks. To further validate the observations on inflammatory immune phenotype of lupus-prone female intestine, immune cells were enriched from colon, cultured for 48 h in the absence of added stimuli, supernatants were tested for the levels of spontaneously secreted cytokines. Figure 1B shows that the amounts of spontaneously released cytokines were significantly higher in females compared to males. Overall, in conjunction with our recent report showing activated T and B cells in the gut mucosa of lupus-prone female mice [27], these observations confirm that pro-inflammatory immune phenotype of gut mucosa in lupus-prone female mice is initiated independent of hormones and at juvenile age.

**Fig. 1:**
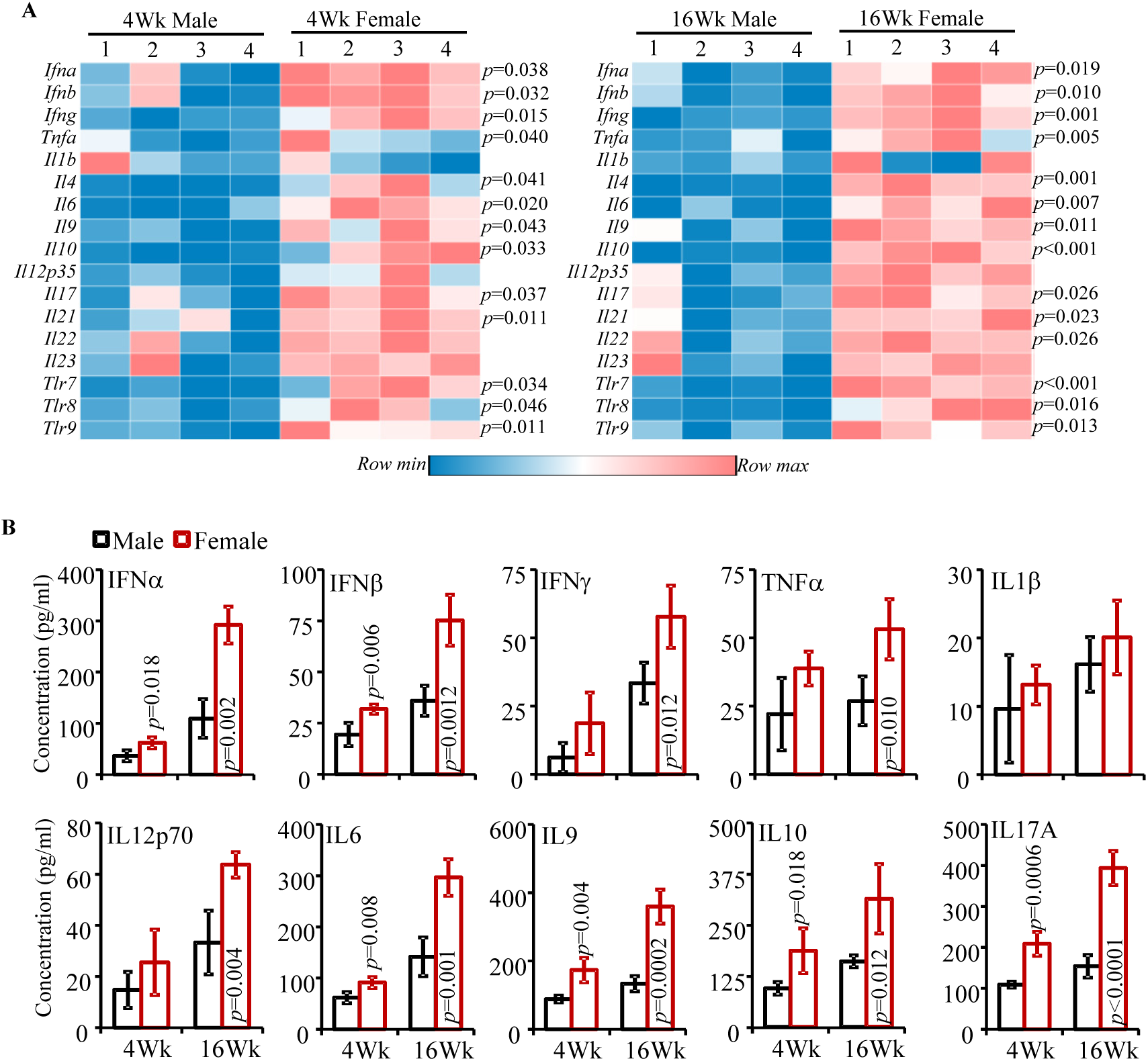
Intestinal immune phenotype in SNF1 mice. A) cDNA preparations from the distal colon of 4 and 16 week old lupus-prone male and female SNF1 mice were subjected to real-time quantitative PCR to assess the expression levels of key cytokines and endosomal TLRs. Expression levels of individual factors were calculated against the β-actin values and values from 4 male and 4 female SNF1 mouse littermates (tested individually) were used for generating the heat-map. Statistical analysis by two-sided *t*-test. Distal ileum from male and female littermates were analyzed similarly and shown in Supplemental Fig. 1. **B)** Immune cells were enriched from collagenase digested single cell suspension of ileum and colon by Percoll gradient centrifugation, equal number of enriched cells (3×10^5^/well in 150 μl medium) from each tissue were cultured for 48 h, and the spent media were subjected to SBA assay to detect the levels of spontaneously released cytokines. Mean±SD values (4 male and 4 female SNF1 mouse littermates; tested individually) are shown. Statistical analysis by two-sided Mann Whitney test. These experiments were repeated at least once using 4 mice/group and showed similar results.

### 4.2. Gut microbiota composition of lupus-prone SNF1 male and female mice is different at adult ages

Differences in the intestinal immune phenotype in male and female SNF1 mice suggested a potential role for gut microbiota in this gender bias. The influence of sex hormone-dependent gut microbiota in T1D gender bias in NOD mice has been reported before [22, 23], however such studies in the context of lupus have not been reported. Our initial studies used age-matched, but not littermate controlled, mice to determine the composition of fecal microbiota in younger (4 week old) and older (24 week old) SNF1 males and females. Principal component (PC) analysis of 16S rRNA gene sequence data of fecal samples from male and female mice showed profound gender specific clustering/separation only at adult age, but not the juvenile group (Supplemental Fig 2A). Accordingly, profound differences in the abundances of microbial communities were observed only in 24 week old male and female mice (not shown). Further, SNF1 females at 24 weeks, the age at which severe nephritis begins to appear, but not 4 week old juvenile females, showed relatively lower gut microbial diversity compared to male counterparts (Supplemental Fig. 2B).

To unambiguously validate these observations, fecal pellets from 8 individually housed littermates (4 males and 4 females) of SNF1 mice were collected at 4 weeks and 16 weeks of age and subjected to microbial community profiling. As shown in Fig. 2A, PC analysis found gender specific β-diversity clustering of gut microbiota only with male and female mouse samples collected at 16 weeks of age. Further, as shown in Fig. 2B, α-diversity / species richness was significantly different only in 16 week fecal samples. Compilation of OTUs to different taxonomical levels showed that fecal microbiota in 16 week old females had significantly lower abundance of Verrucomicrobia phyla members and is relatively richer, albeit not significant statistically, in Bacteroidetes phyla members (Fig. 2C and Supplemental Fig. 3). Interestingly, a significant increase in Bacteroidetes abundance was observed in females at 16 weeks of age compared to 4 weeks (Supplemental Fig. 4A). Since Fig. 2A showed gender dependent clustering only with samples from 16 week old mice, these samples were compared for abundances of microbial communities at the genus level. As observed in Fig. 2D, compared to male littermates, samples from females at 16 weeks of age had profoundly higher abundance of Bacteroides and Parabacteroides genus members, and lower Dysgonomonas genus members of Bacteroidetes phylum. Further, significant differences in large number of minor communities were observed between 16 week fecal samples of males and females. As observed in Supplemental Fig. 4B, significant age-dependent differences in the abundance of multiple microbial communities were observed in both males and females when samples collected at 4 weeks were compared to that of 16 weeks of age.

**Fig. 2:**
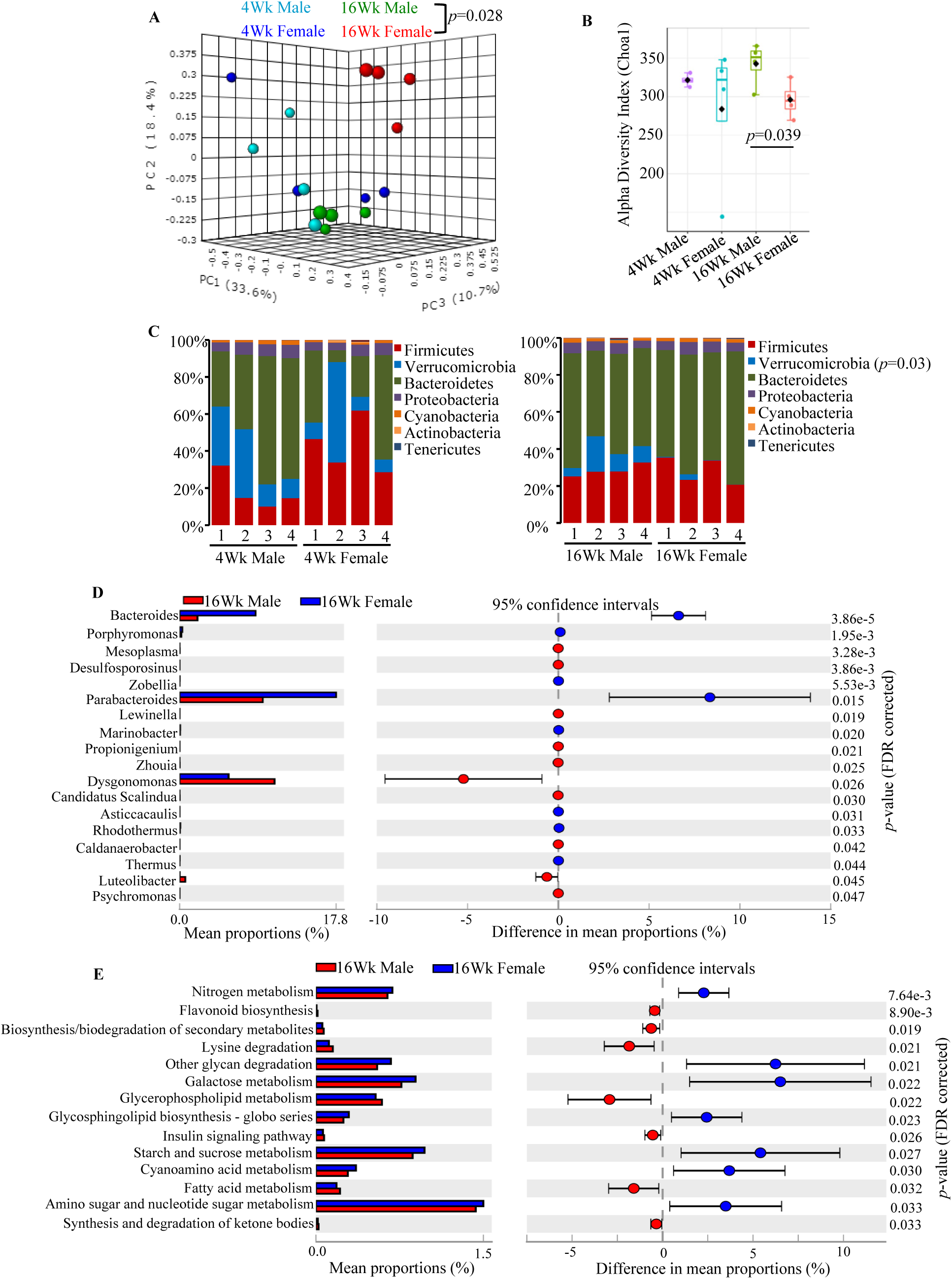
Gut microbiota composition in lupus-prone SNF1 male and female mice. Four male and 4 female SNF1 mouse littermates were individually housed upon weaning at 3 weeks of age, fresh fecal pellets were collected at 4 and 16 weeks of age, DNA preparations were subjected to 16S rRNA gene (V3/V4 region)-targeted sequencing using the Illumina MiSeq platform. The OTUs that were compiled to different taxonomical level based upon the percentage identity to reference sequences (i.e. >97% identity) and the percentage values of sequences within each sample that map to specific phylum and genus were calculated employing 16S Metagenomics application of Illumina BaseSpace hub. OTU.Biom tables were also generated from the sequencing data employing QIIME preprocessing application of BaseSpace. **A)** OTU.Biom table data were used for generating principal component analysis (PCA) plots of samples representing β-diversity (Bray Curtis distance). **B)** α-diversity (Chao1)/species richness comparison of all 4 groups. **C)** Relative abundances of 16S rRNA gene sequences in fecal samples of individual mice at phyla level. **D)** Mean relative abundances of sequences representing major and minor microbial communities (>0.1 % of total sequences) at genus level. **E)** Normalized OTU.biom table was used for predicting gene functional categories using PICRUSt application and selected level 3 categories of KEGG pathway with statistically significant differences (between 16 week males and 16 week females) are shown. Statistical analyses were done employing two-sided Welch’s *t-test* and the *p*-values were FDR corrected using Benjamini and Hochberg approach. Additional analyses of this sequencing data are shown in Supplemental Figs. 3 & 4.

We then employed PICRUSt application for OTUs based *in silico* prediction of metabolic functions that may be associated with intestinal pro-inflammatory immune phenotype and the disease process in lupus-prone females. The PICRUSt analysis of data from 16-week old males and females showed that a variety of metabolic pathways such as glycan degradation and glycosphingolipid biosynthesis pathways, and nitrogen, amino sugar, nucleotide sugar, galactose, starch and sucrose metabolism pathways are overrepresented in females at 16 weeks of age (Fig. 2E). On the other hand, pathways involved in secondary metabolite synthesis and degradation, fatty acid, glycerophospholipid and lysine metabolism are underrepresented in 16 week old females compared to male counterparts. Although further analysis and additional studies are needed to understand if these metabolic pathways contribute to modulation of intestinal immune response and disease progression, these observations further support the notion that microbial communities are functionally different in lupus-prone adult males and females.

### 4.3. Gut microbiota depletion suppresses autoimmune progression in female SNF1 mice

Although we observed a pro-inflammatory immune phenotype in female SNF1 mouse intestine as early as juvenile age, significant differences in the overall gut microbiota composition between males and females was detected only at adult ages. Nevertheless, we examined if microbiota depletion impacts systemic autoimmunity and disease incidence in these mice. Both male and female SNF1 mice were given a broad-spectrum antibiotic cocktail in drinking water, starting immediately after weaning (3 weeks of age) to deplete the gut microbiota (Supplemental Fig. 5) and monitored for proteinuria and serum autoantibody levels. As shown in Fig. 3A, compared to mice with intact gut microbiota, severe nephritis was significantly delayed in majority of the female SNF1 mice that were given antibiotics. While about 80% of control females developed severe nephritis by 31 weeks of age, only about 40% of the microbiota depleted mice showed similar degree of disease. Intriguingly, microbiota depletion had no impact on the disease progression in male SNF1 mice. Fig. 3B which shows differences in the degree of proteinuria in antibiotic treated and control mice further supports the observation that the overall disease progression was suppressed significantly in microbiota depleted females, but not in male SNF1 mice. In addition, determination of serum autoantibodies against dsDNA and nucleohistone revealed significantly lower autoantibody production in microbiota-depleted female SNF1 mice compared to control mice (Fig. 3C). On the other hand, autoantibody responses were comparable in microbiota depleted and control male SNF1 mice.

**Fig. 3:**
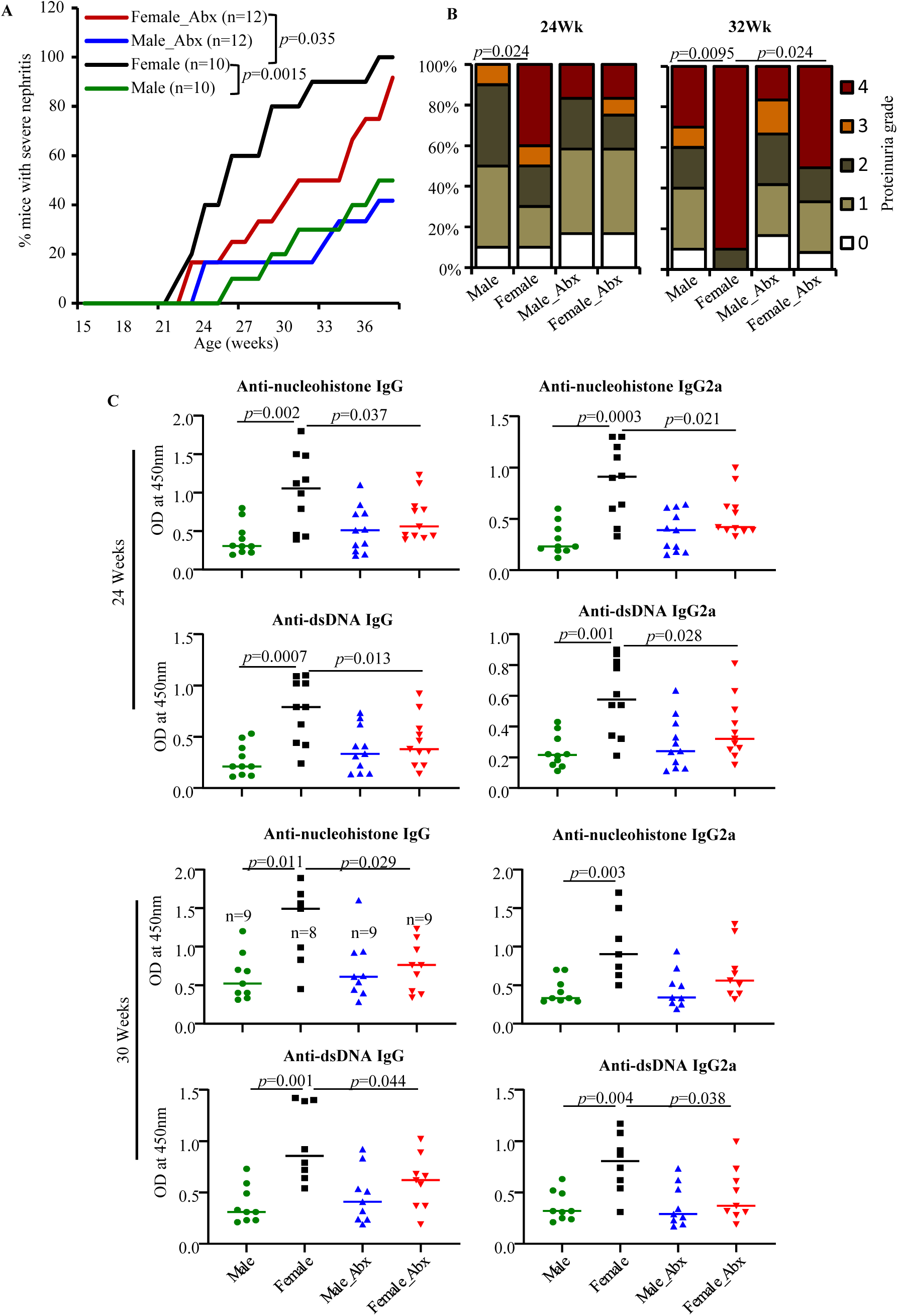
Impact of gut microbiota depletion on autoimmune progression. Male and female SNF1 mice were maintained on regular drinking water or drinking water containing broad-spectrum antibiotic cocktail starting at 3 weeks of age. Urine and serum samples collected from individual mice at different time-points were examined for proteinuria and auto-antibody levels. **A)** Protein levels in urine samples were quantified by Bradford assay. Percentage of mice with severe nephritis as indicated by high proteinuria (≥5mg/ml) is shown. *p-*values by log-rank test. **B)** Severity of proteinuria/nephritis in different groups of mice at different time points is shown. Nephritis severity was scored based on urinary protein levels as follows; 0: 0-1mg/ml, 1:1-2mg/ml, 2:2-5 mg/ml, 3: 5-10mg/ml and 4: ≥ 10mg/ml. *p*-values by 2-sided Chi-Square test. **C)** Serum levels of total IgG and IgG2a antibodies specific to nucleohistone and dsDNA were assessed by ELISA for samples collected at 24 and 30 weeks of age. Mean ±SD of OD values of samples (1:1000 dilution) from 10 mice/group for controls and 12 mice/group for antibiotic treated mice are shown. *p*-values by two-sided Mann Whitney test.

### 4.4. Microbiota depletion suppresses intestinal pro-inflammatory immune phenotype in female SNF1 mice

Since depletion of gut microbiota resulted in slower autoimmune progression in females, intestinal immune phenotype of mice with intact and depleted gut microbiota was determined in cohorts of young (5 weeks; pre-puberty) and adult (16 week) mice. As observed in Fig. 4A, the expression levels of pro-inflammatory cytokines in the colon of microbiota depleted females were significantly lower compared to control females, irrespective of the age. On the other hand, expression of *Il10*, an immune regulatory cytokine, was not diminished in microbiota depleted younger mice and significantly higher in microbiota depleted older mice compared to their counterparts with intact microbiota. Further, the levels of cytokines secreted by intestinal immune cells (Fig. 4B) showed similar trend with that of mRNA levels. Interestingly, *Tlr7, Tlr8, and Tlr9* expression levels in the colon of microbiota depleted females were not significantly different from that of control mice. Of note, in agreement with proteinuria and autoantibody levels shown in Fig. 3, expression levels of pro-inflammatory cytokines in the colon were not significantly different in microbiota depleted and control SNF1 males (not shown).

**Fig. 4:**
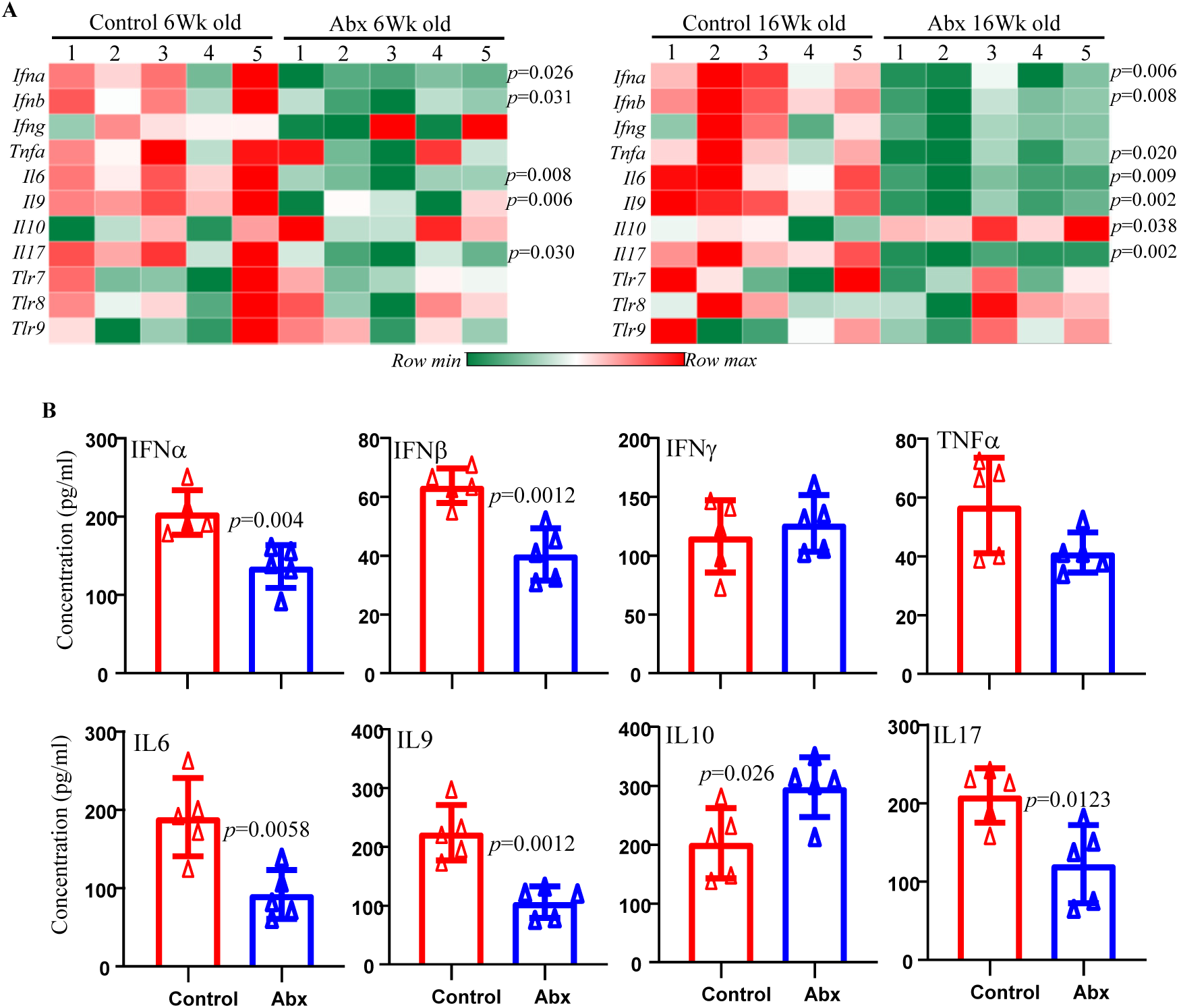
Impact of microbiota depletion on intestinal pro-inflammatory immune phenotype. A) cDNA preparations from the distal colon of 5 week and 16 week old control and antibiotic cocktail (Abx) treated female SNF1 mice were subjected to real-time quantitative PCR to assess the expression levels of key cytokines and endosomal TLRs. Relative expression levels of individual factors were calculated against the β-actin values and values from a total of 5 control and 5 antibiotic treated females (tested individually) were used for generating the heat-map. **B)** Immune cells were enriched from collagenase digested single cell suspension of whole ileum and colon of 16 week old mice (4 mice/group) by Percoll gradient centrifugation, equal number of cells (3×10^5^/well in 150 μl medium) from each tissue were cultured for 48 h and the spent media were subjected to SBA assay to detect spontaneously released cytokines. Statistical analyses by two-sided Mann Whitney test. These experiments were repeated at least once using 3 mice/group and similar results were obtained.

### 4.5. Castration of male SNF1 mice causes changes in gut microbiota composition

Our observations (Supplemental Fig. 4) show that both male and female SNF1 mice present age dependent changes in gut microbiota composition. However, age-dependent β-diversity clustering (Fig. 2A) and impact of microbiota depletion (Figs. 3 & 4) were observed only with females. This suggested that the gut microbes contribute to higher disease incidence in females. Of note, previous reports have shown androgen dependent changes in the gut microbiota in NOD mouse model of T1D [22, 23] and resistance to disease compared to female counterparts. Therefore, we examined if androgen has an influence on the gut microbiota in lupus-prone male SNF1 mice. Juvenile mice were castrated to curtail androgen production (Supplemental Fig. 6), and examined for changes in fecal microbiota composition. PC analysis graphs (Fig. 5A) show that samples from castrated and mock-surgery control mice segregated better at an adult age (post-castration) compared to juvenile age (pre-castration). Although the difference in microbial alpha diversity indices was not statistically significant in castrated and control mice (Fig. 5B), lower abundances of Bacteroidetes and Verrucomicrobia and higher abundances of Proteobacteria and Firmicutes phyla were observed in castrated males compared to controls (Fig. 5C). Figure 5D shows that the abundance of multiple minor microbial communities (those with <1% of all identified sequences) at genus level were significantly different in castrated and non-castrated SNF1 males. Overall, these observations show that gut microbiota compositions are different in castrated and non-castrated male SNF1 mice and, as reported by others in NOD mice [22, 23], testosterone may be contributing to these differences.

**Fig. 5:**
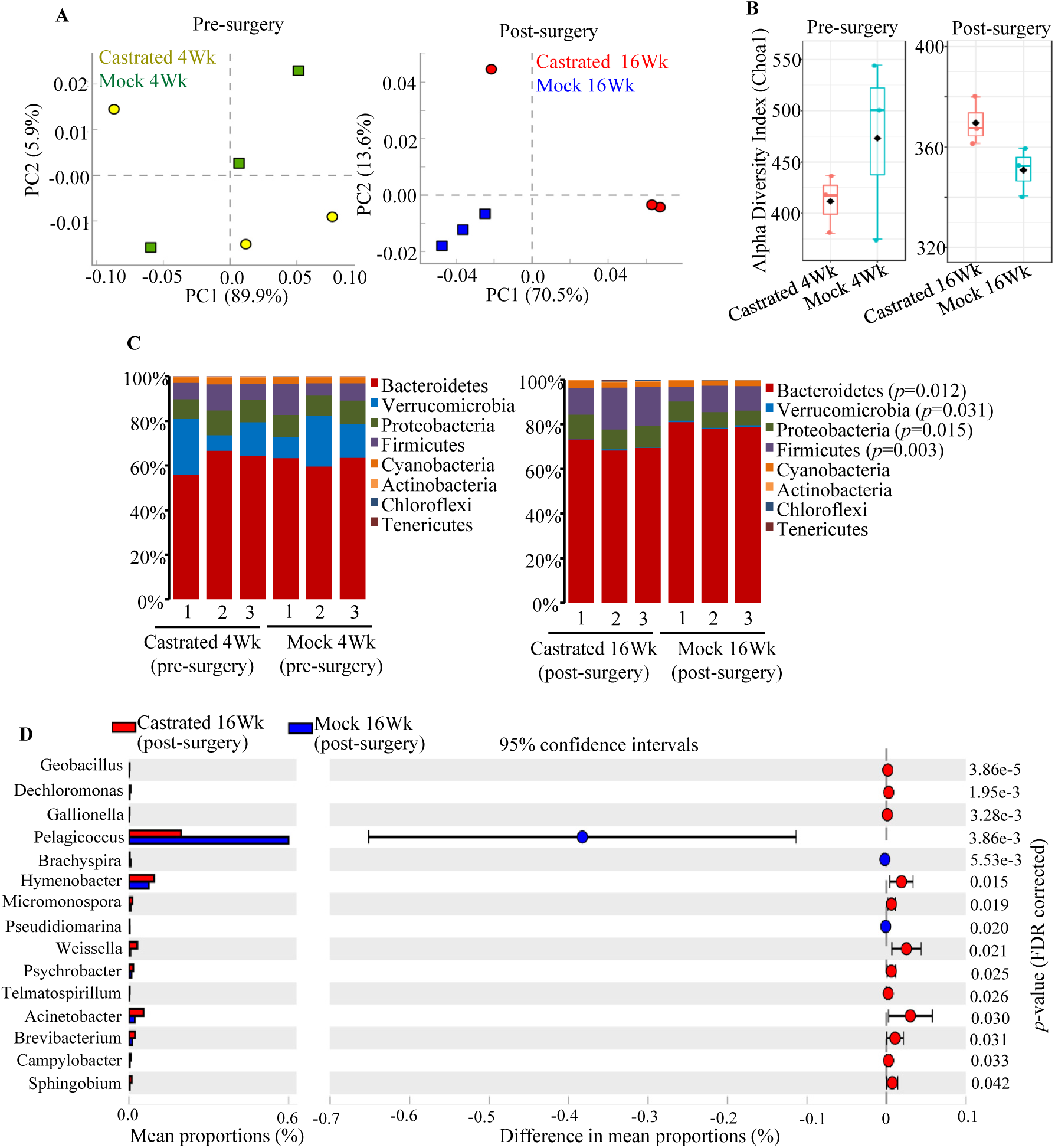
Impact of castration of SNF1 males on gut microbiota. Four week old SNF1 males were subjected to orchidectomy or mock-surgery (3 mice/group), fecal pellets were collected from individual mice before the surgery (pre-surgery) and at 20 weeks of age (post-surgery). DNA prepared from these fecal samples were subjected to 16S rRNA gene-targeted sequencing and analysis as described for Fig. 2. **A)** OTU.Biom table data were used for generating principal component analysis (PCA) plots. **B)** Relative abundances of 16S rRNA gene sequences in fecal samples of individual mice collected pre-and post-surgery at phyla level. **C**) Mean relative abundances of sequences compiled to genus level in post-surgery (20 week) samples are shown. Statistical analyses were done employing two-sided Welch’s *t*-test and FDR corrected using Benjamini and Hochberg approach.

### 4.6. Castration of male SNF1 mice results in modestly accelerated autoimmune progression

Since depletion of gut microbiota had no impact on autoimmune progression in male SNF1 mice (Fig. 3) and castration appeared to influence the gut microbiota composition in them (Fig. 5), we examined if autoimmune progression and disease incidence in SNF1 males are impacted by low testosterone levels resulting from castration. Castrated and mock-surgery control male mice were monitored for proteinuria for up to 36 weeks of age. As observed in Fig. 6, although not significant statistically, castrated males showed relatively earlier onset and higher incidence of disease, higher severity of nephritis, and higher anti-nucleohistone and anti-dsDNA antibody responses. Of note, overall immune phenotype of gut mucosa was not significantly different in castrated and control mice (not shown). These results along with the observation that castrated and non-castrated mice show differences in gut microbiota suggested that androgen may directly or indirectly (through gut microbiota) contribute to the lower susceptibility of male SNF1 mice to systemic autoimmunity.

Since microbiota depletion did not impact disease incidence in male SNF1 mice (Fig. 3), microbiota transfer experiment was carried out to determine if androgen influenced microbiota can influence the disease progression. Puberty aged SNF1 females received cecum microbial preparations from castrated males, mock surgery control males and age matched females (20 weeks of age) by oral gavage. These donor male mice had undergone castration or mock surgery at juvenile age and produced little or no testosterone (Supplemental Fig. 6). As shown in Fig. 6D, female SNF1 mice that received cecum microbial preparations from all three adult groups showed comparable disease progression and proteinuria severity. Further, the levels of anti-dsDNA and anti-nucleohistone antibody levels were comparable in these groups of mice (Supplemental Fig. 7). Of note, while all microbiota recipient groups of mice showed relatively slower disease progression compared to that of non-recipient control, mice that received androgen shaped gut microbiota failed to show better protection of mice from lupus. These results, along with the observation that microbiota depletion has no impact on lupus associated features in males (Fig 3), suggest that androgen mediated protection of male SNF1 mice from lupus is not microbiota dependent.

**Fig. 6:**
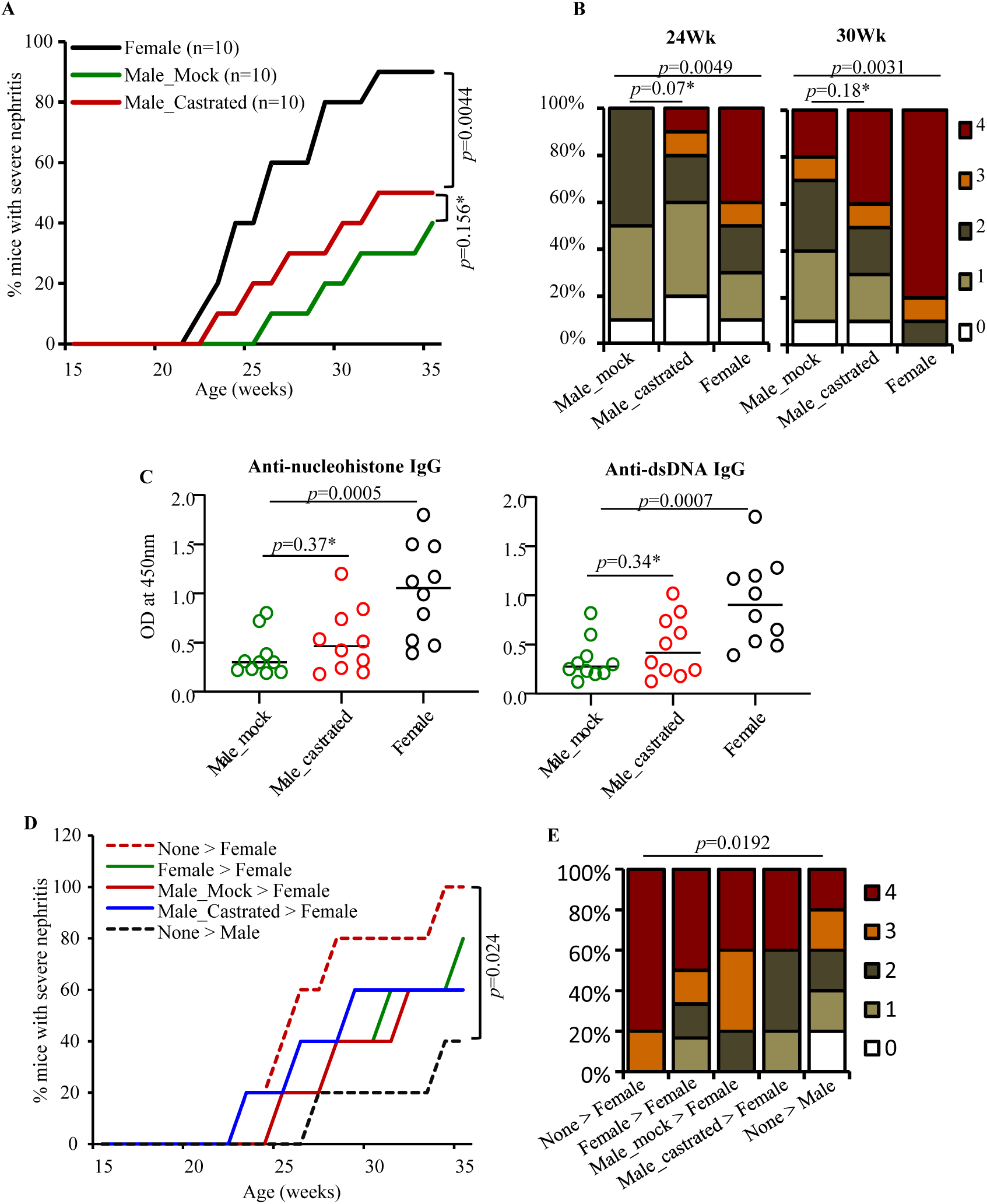
Impact of Castration of SNF1 males on lupus autoimmunity. A) Four week old male SNF1 littermates were subjected to orchidectomy or mock-surgery (10 mice/group) and monitored for disease progression as detailed for Fig. 3. **A)** Protein levels in urine samples were quantified by Bradford assay. Percentage of mice with severe nephritis as indicated by high proteinuria. *p-*values by log-rank test. **B)** Severity of nephritis in different groups of mice at different time points. Nephritis severity was scored based on urinary protein levels. *p*-values by 2-sided Chi-Square test. **C)** Serum levels of IgG antibodies against nucleohistone and dsDNA were assessed by ELISA using samples collected at 28 weeks of age. Mean ±SD of OD values of samples from 10 mice/group. *p*-values by two-sided Mann Whitney test. **D)** Cecum microbial preparations from 20 week old indicated groups of mice were given to pre-pubescent female SNF1 mice (5 recipients or control mice/group) and monitored for disease progression, and percentage of mice with severe nephritis as indicated by high proteinuria is shown. *p-*values by log-rank test. **E)** Severity of nephritis in different groups of mice at 28 weeks of age is shown. *p*-values by 2-sided Chi-Square test. *not statistically significant. Serum levels of IgG antibodies against nucleohistone and dsDNA assessed by ELISA are shown in supplemental Fig. 7.

## 5. Discussion

Gender bias is prevalent in many major autoimmune diseases including SLE [41–43]. The influence of sex hormones on lupus autoimmunity in general and gender bias in particular has been extensively studied. While estrogen has been linked to both cause and pathology of the disease [44–46], testosterone has generally been found to have a protective effect [46, 47]. Interestingly, recent studies using SPF and germ-free (GF) NOD mice have linked the sex-hormone, primarily androgen,-influenced gut microbes and their metabolites to gender bias in autoimmunity [22, 23]. It has been shown that, similar to T1D mouse model, SLE mouse models in GF background do develop disease, indicating that gut microbes are not required for systemic autoimmune response in them [24, 25]. However, the role of gut microbiota in gender bias associated with lupus has not yet been fully realized. In this regard, our recent report [27] showed, for the first time, that the immune phenotype of gut mucosa is significantly different in lupus-prone male and female SNF1 mice. Pro-inflammatory immune phenotype of gut mucosa appears in female mice at juvenile age, much before the production of sex hormones. Further, gut mucosa of female SNF1 mice carry higher number of plasma cells compared to their male counterparts even at juvenile age. These observations indicated that immune response initiated in the gut mucosa may be involved in the initiation and perpetuation of systemic autoimmunity and determining lupus associated gender bias. In the current study, we found significant influence of gut microbiota on intestinal pro-inflammatory immune phenotype and systemic autoimmune progression in females, but not in males. Further, we show that while androgen has an influence on gut microbiota composition in lupus model as reported in T1D model by others [22, 23], it appears to promote microbiota-independent modest protection from lupus autoimmunity.

Our observation that gut microbiota compositions of male and female lupus-prone mice differ significantly only at adult ages is in agreement with previous reports in other models [22, 23, 48, 49]. It has been suggested that maternal microbiota acquired at young ages could be influenced by the host factors such as hormones and environmental factors and eventually change/mature. In a disease-prone background, these changes could profoundly influence the disease outcome [10, 14, 15, 23, 50]. The disease susceptibility and changes/differences in the abundances of microbial communities, including increase in Bacteroidetes abundance and diminished diversity close to disease onset, in lupus-prone males and females are in agreement with previous reports [26, 32, 51]. Interestingly, in contrast to the observations in NOD mouse model where the age dependent drift in microbial composition was prominent in males [22, 23], lupus-prone SNF mice showed more pronounced changes in females compared to male littermates, implying a role for gut microbes in the pro-inflammatory immune phenotype and higher disease incidence in these females.

Profiling of intestines of lupus-prone male and female SNF1 mice for various markers demonstrated that both colon and ileum of females express higher levels of pro-inflammatory cytokines as well as TLRs as early as juvenile age. Endosome TLRs such as TLR7, TLR8 and TLR9 are widely implicated in lupus [36–39] and many of the therapeutic agents that are already being used in the clinic do target the endosome pathway [52]. Pro-inflammatory cytokine factors such as IFN-α, IFN-β, IL-17 and IL9 have also been linked to pathogenesis associated with SLE [36, 39, 53–56]. Importantly, signaling through endosomal TLRs triggers production of pro-inflammatory cytokines such as type 1 IFNs [39, 57, 58]. Compared to female SNF1 mice with intact gut microbiota, lower expression of only pro-inflammatory cytokines, but not the immune regulatory cytokine IL10 or endosomal TLRs, were observed in pre-puberty and adult mice upon depletion of gut microbiota. Moreover, microbiota depletion associated suppression of pro-inflammatory cytokines in the intestinal mucosa occurs mainly in females, but not in males. This is intriguing because the composition of gut microbiota in male and female mice, as reported by others in the past [24, 25], are comparable at younger ages. Nevertheless, these microbiota dependent features of gut mucosa of male and female SNF1 mice explain why systemic autoimmunity and the disease incidence are significantly suppressed only in females upon microbiota depletion. Of note, although previous studies using lupus-prone GF mice did not conclusively demonstrate a role for gut microbiota in autoantibody production and disease incidence [24, 25], as observed upon selective depletion of gut microbiota using selected antibiotics in MRL/lpr mice [59], we found that treatment using broad-spectrum antibiotic cocktail suppresses autoimmunity in SNF1 females.

Our observations, consistent with previous reports [24, 25, 48, 49], that gut microbiota compositions in SNF1 male and female littermates are significantly different only at adult age, but not juvenile age, do argue against the notion that differences in the composition of gut microbiota could be responsible for the pro-inflammatory immune phenotype of juvenile SNF1 female intestine. Nevertheless, microbiota depletion did result in diminished expression of pro-inflammatory cytokines in the intestine of not only adult, but also the juvenile, SNF1 females. Although studies using GF lupus-prone mice are needed to further evaluate the true impact of microbiota on intestinal immune phenotype, our observation raises the possibility that the magnitude/dosage of microbiota-gut mucosa interaction, rather than the microbiota composition, is different in male and female SNF1 mice. Higher expression levels of microbiota interacting receptors in the gut mucosa could contribute to higher expression of pro-inflammatory factors in the female intestine and the disease associated gender bias. A likely scenario, based on our qPCR data, is that higher levels of TLRs, such as TLR7, TLR8 and TLR9, in the gut mucosa of lupus-prone females could facilitate higher dose interaction with gut microbiota, resulting in more pronounced inflammatory cytokine response. In this regard, previous studies have shown that TLR7 and TLR8 stimulation promotes significantly higher pro-inflammatory cytokine response in females than in males [36–39]. A recent study using mouse models has shown that dosage of X-linked TLR8 plays a major role in the higher incidence of SLE in females [39]. Another study using samples from lupus patients has demonstrated that TLR7 ligands induce higher IFN-α production in females [36]. Of note, the fact that the expression of pro-inflammatory cytokines is much higher in adult SNF females compared to prepubescent mice suggest a progressive increase in the recruitment and/or activation of immune cells in the gut mucosa causing a vicious cycle of pro-inflammatory responses contributing to eventual systemic autoimmunity.

Our observation that significant impact of microbiota depletion on gut immune phenotype and autoimmune progression is observed only in female, but not in male, SNF1 mice is also intriguing. One possible explanation is that under normal circumstances, unlike in lupus-prone females, low dose gut mucosa-microbiota interaction produces only a modest cytokine response in control mice with intact microbiota and, as reported by others [46, 47], testosterone alone may have a disease protective effect. Previous reports have shown that testosterone influenced gut microbiota, rather than the higher testosterone levels, protects NOD mice from T1D [22, 23]. In agreement with these reports, castration of male SNF1 mice did cause changes in the overall composition of gut microbiota, primarily by altering the abundances of minor communities. However, our microbiota transplant studies showing lack of superior protection of female mice by androgen-influenced gut microbiota, compared to microbiota from females and castrated males, suggest that modest increase in disease susceptibility in castrated male SNF1 mice could be due to the lack of testosterone, but may not be microbiota dependent. This notion has been further validated by our observation that microbiota depletion has no impact on autoimmunity and disease incidence in male SNF1 mice. In fact, although impact on gut microbiota was not assessed, previous reports have shown that androgen contributes to slower disease progression and lower lupus-like incidence lupus-prone mice [47, 60, 61]

Overall, our study, for the first time, demonstrates distinct roles for gut microbiota in lupus-prone female and male mice contributing to gender bias in the disease outcomes. In females, gut microbiota appears to have composition-independent, gut mucosa interaction-dose dependent, role in promoting intestinal pro-inflammatory immune response and eventual systemic autoimmunity. On the other hand, androgen contributes to protection of male SNF1 mice from the disease microbiota independently. These observations also suggest that the mechanisms associated with gender bias in lupus model may be different from that of T1D model, which is primarily due to protection of males from the disease by testosterone-influenced gut microbiota [22, 23]. Hence, comprehensive studies using GF mice are needed in the future to delineate the microbiota-dependent and-independent mechanisms of gender bias in lupus autoimmunity.

## Abbreviations

SLE: systemic lupus erythematosus
SNF1 mice: (SWRxNZB)-F1 mice
T1D: Type 1 Diabetes Mellitus
EAE: experimental autoimmune encephalitis
nAg: nuclear antigen
PRR: pattern recognition receptor
PP: Peyers’ patch
MLN: mesenteric lymph node
LP: Lamina Propria
SI: small intestine
Treg: regulatory T cells
GF: germ-free
TLR: toll-like receptors
EBV: Epstein Barr virus
RA: rheumatoid arthritis

## Acknowledgments

This work was supported by internal funds from MUSC, National Institutes of Health (NIH) grants R21AI136339, R01AI138511 and R01AI073858, and Lupus research Institute (LRI) award to C.V. and P60AR062755 (NIH/NIAMS) to G.G. through a pilot grant to C.V. B.M.J. performed the experiments and reviewed the paper, M.C.G. performed the experiments and reviewed the paper, R.B. performed the experiments, R.G. performed experiments and reviewed the paper, G.G. reviewed the paper, and C.V. designed the study, performed the experiments, and wrote the paper. Dr. Vasu is the guarantor of this work and, as such, had full access to all the data in the study and takes responsibility for the integrity and accuracy of the data and analysis. The authors are thankful to MUSC’s DLAR Veterinary staff for mouse castration service. The authors are also thankful to Cell and Molecular Imaging, Pathology, Proteomics, immune monitoring and discovery, and flow cytometry cores of MUSC for the histology service, microscopy, real-time PCR, FACS and multiplex assay instrumentation support. We also thank Drs. Raad Z Gharaibeh, Cory R. Brouwer, and Anthony A Fodor, Department of Bioinformatics and Genomics, UNC Charlotte for the initial guidance on analysis of MiSeq data and valuable comments and suggestions.

