## Supplemental data for "Gut microbiota differently contributes to intestinal immune phenotype and systemic autoimmune progression in female and male lupus-prone mice"

Supplemental Figure 1

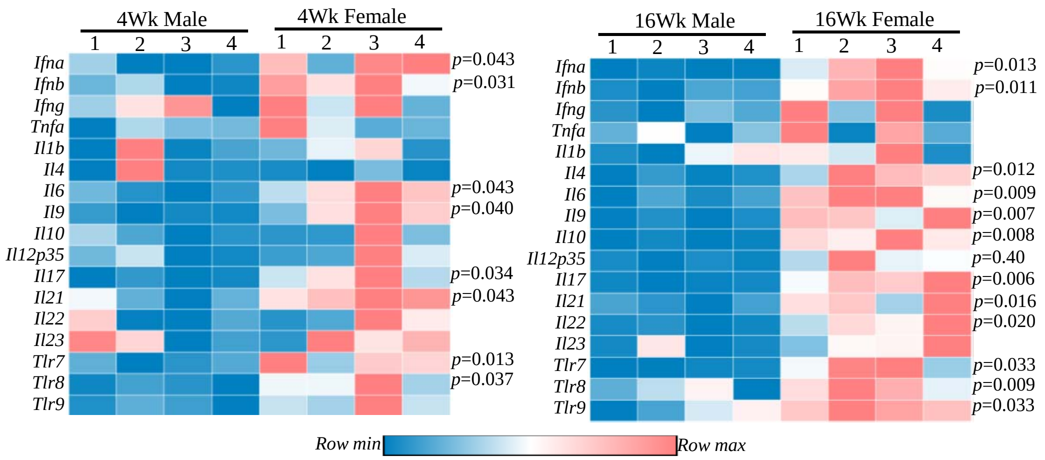


*Supplemental Fig. 1:* Real-time quantitative PCR to assess the expression levels of key cytokines and endosomal TLRs was performed using cDNA prepared from the distal ileum tissues of mice described under Fig. 1. Statistical analysis by two-sided *t*-test.

### Supplemental Figure 2

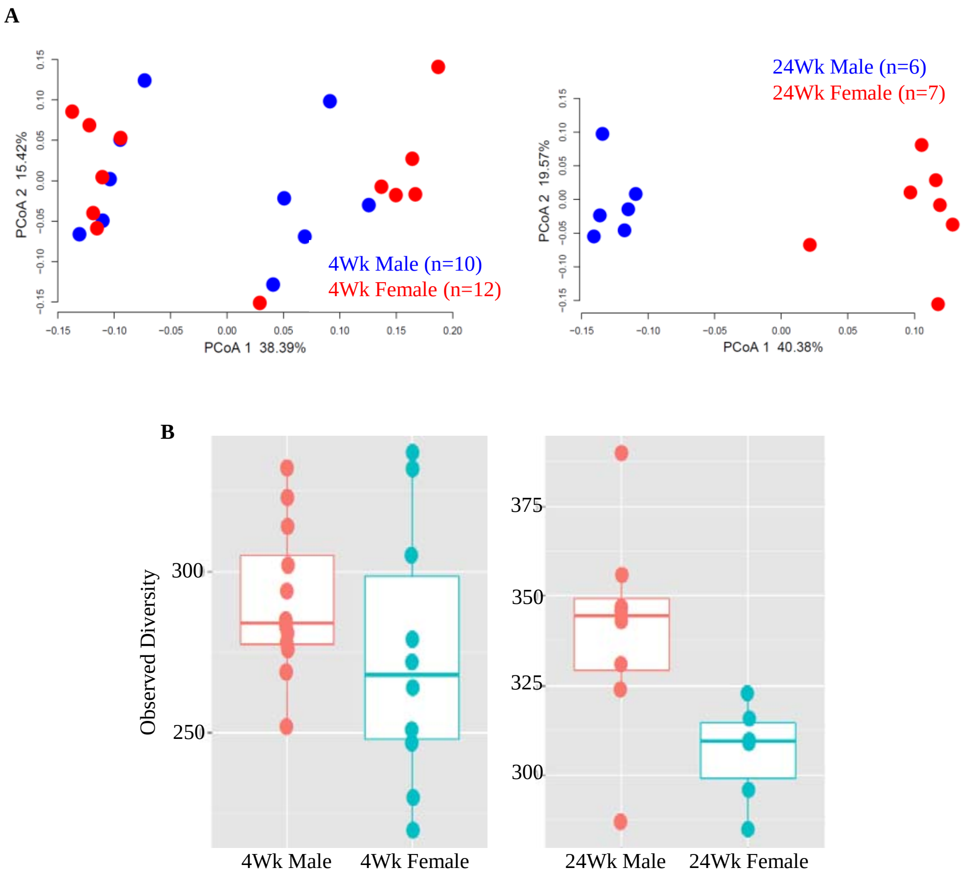


*Supplemental Fig. 2:* DNA prepared from the fecal samples collected of 4 week old and 24 week old male and female SNF1 mice were subjected to 16S rRNA gene sequencing and analyzed as described under material and methods. Alpha (**A**: Bray Curtis distance) and Beta diversity (**B**: species richness) measures are shown.

### Supplemental Figure 3

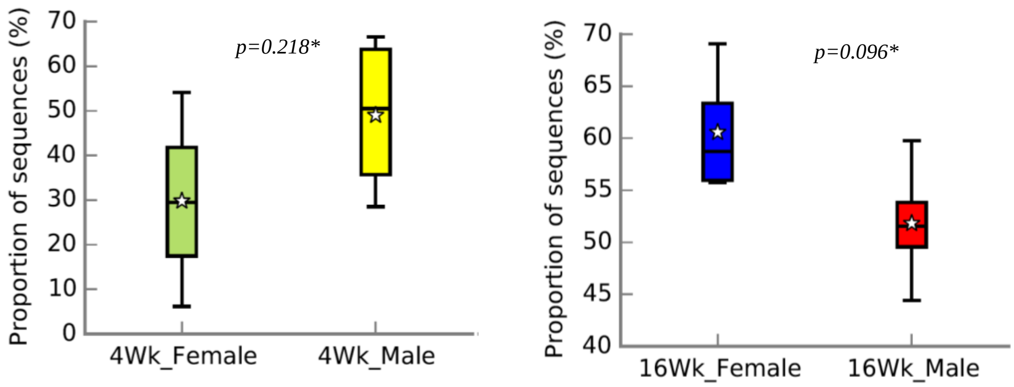


*Supplemental Fig. 3:* DNA prepared from the fecal samples collected from individually housed SNF1 male and female littermates at 4 and 16 weeks of age were subjected to 16S rRNA gene sequencing and analyzed as described in Fig.2. Abundances of Bacteroidetes phylum in different groups of mice are shown. *not statistically significant.

### Supplemental Figure 4

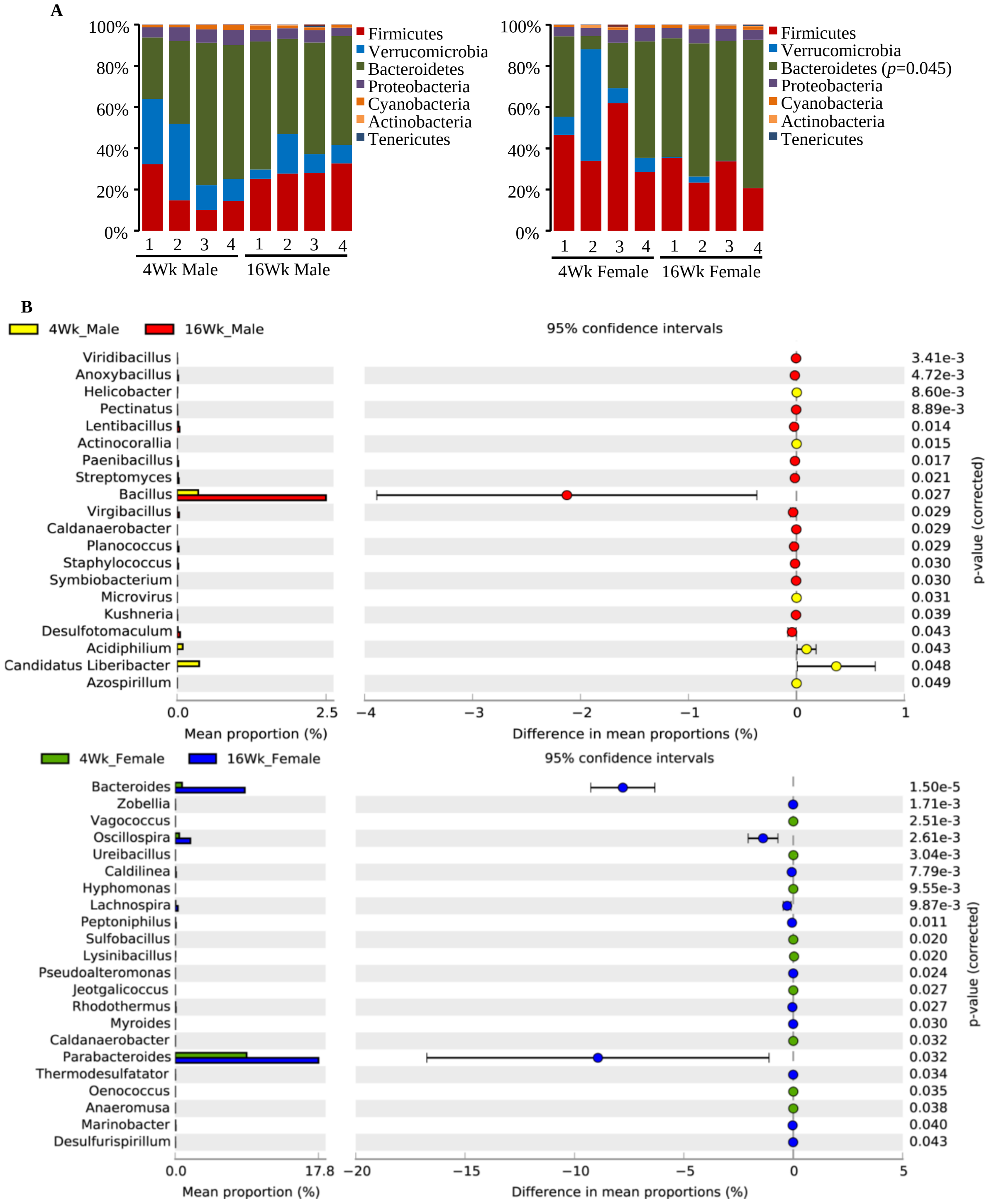


*Supplemental Fig. 4:* DNA prepared from the fecal samples collected from individually housed SNF1 male and female littermates at 4 and 16 weeks of age were subjected to 16S rRNA gene sequencing and analyzed as described in Fig.2. Figure 2 shows key analyses of the same sequences. Additional comparisons of microbial community profiles at phyla level **(A)** and genus level **(B)** in samples collected from the same gender at 4 and 16 week time-points are shown here.

### Supplemental Figure 5

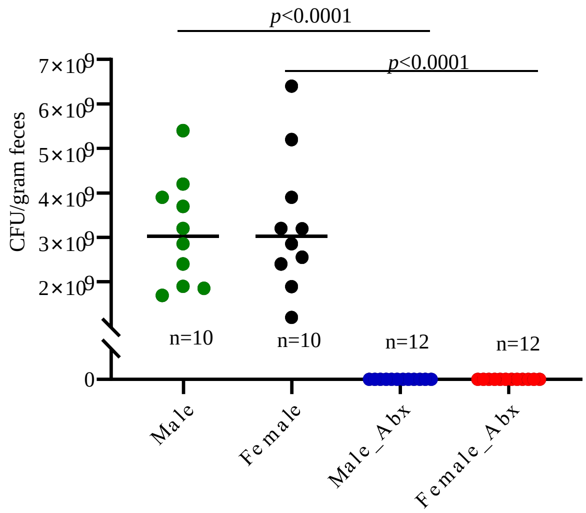


*Supplemental Fig 5:* Depletion of gut microbiota using antibiotics. Male and female SNF1 mice were given a broad-spectrum antibiotic cocktail (ampicillin (1 g/l), vancomycin (0.5 g/l), neomycin (1 g/l), and metronidazole (1 g/l) -containing drinking water starting at juvenile age as described for Fig. 3. Fecal pellets collected from individual test and untreated controls at 8 weeks of age were suspended and diluted in sterile PBS, plated onto BHI medium plates under anaerobic and aerobic conditions for up to 72 h, the total number of colonies were counted, and colony forming units (CFU)/gram initial fecal material were calculated. n= 10 mice/group. *p*-values by two-sided Mann Whitney test.

Supplemental Figure 6

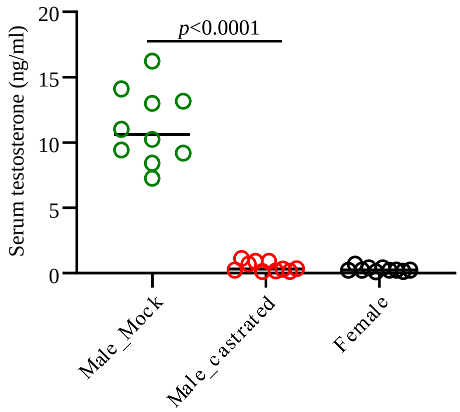


*Supplemental Fig 6: Serum testosterone levels in castrated and control mice.* Four week old male SNF1 mice were subjected to orchidectomy or mock-surgery (10 mice/group) as described in Fig. 5 and serum levels of testosterone were measured by ELISA at week 12 of age. Age matched females were included as additional controls. *p*-values by two-sided Mann Whitney test.

### Supplemental Figure 7

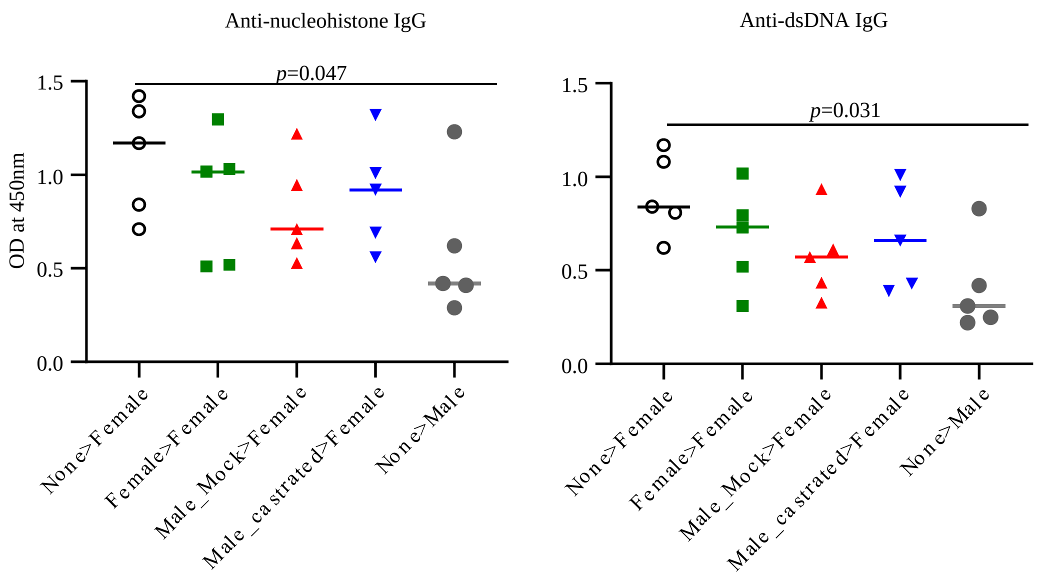


*Supplemental Fig. 7: Autoantibody levels in fecal microbiota recipient mice.* Cecum microbial preparations from indicated groups of mice were given to 6 week old female SNF1 mice (5 recipients or control mice/group) as described in Fig. 6D and 6E. Serum levels of IgG antibodies against nucleohistone and dsDNA were assessed by ELISA using samples collected at 28 weeks of age. Mean ±SD of OD values of samples from 5 mice/group. *p*-values by two-sided Mann Whitney test.
